# Live imaging of bacterial actin MreBs from *Spiroplasma* causing helicity switching of a minimal synthetic cell

**DOI:** 10.1101/2025.05.23.655722

**Authors:** Yoshiki Tanaka, Hana Kiyama, Yusuke V. Morimoto, Makoto Miyata

## Abstract

*Spiroplasma* swim by switching their helical body into right- and left-handed. Helicity formation and switching can be reconstituted in an immotile minimal synthetic bacterium, JCVI-syn3B by introducing the pair of bacterial actin, MreB4 and MreB5. Here, we analyzed MreB behaviors optically, to investigate this unknown mechanism. We tried MreB4 fluorescence labeling by protein fusion. The labeling was not achieved, because the fusion of fluorescent protein or peptide to 16 positions resulted in immotile constructs. These results may suggest that MreB4 has many interaction interfaces with other proteins. Induced expression of MreB4 to cells with constitutive MreB5 expression resulted in earlier and higher frequency of motile cells, distinct from the results of constitutive MreB4 and inducible MreB5. Next, the behavior of labeled MreB5 was analyzed. Photobleaching and photoactivation suggested static behavior of MreB5 during cell movements. Cell treatment by A22, a MreB polymerization inhibitor caused helix deformation, movement stall, and diffusion of MreB5 fluorescence, suggesting that A22 sensitive MreB5 interaction should be involved in helix formation and motility.

**Significance:** *Spiroplasma* MreB is a unique bacterial actin that causes motility with a different mechanism from other actins. Here, we investigated the helicity switching involved in the motility. Fluorescently traced MreB behaviors suggested that the motility mechanism is not coupled with its replacement in the internal structure.

## Introduction

*Spiroplasma. eriocheiris*, a crustaceans parasite [1] belongs to wall-less class Mollicutes bacteria [2,3]. It swims by switching its helical handedness between right and left. The boundary of the handedness called “kink” propagates along the axis, rotating the cell body to push water backward. They have no structural similarities with the machineries of Spirochetes helix rotation [4-6]. *Spiroplasma* motility machinery is caused by ribbon-like structure positioning along the innermost line of the helix [5,7]. The structure is composed of the *Spiroplasma*-specific protein Fibril, and five actin homologs MreB1-5 [7-11]. Previously, our group transferred genes of these proteins to a minimal synthetic bacterium JCVI-syn3B (syn3) [12,13], which has minimal set of genes for reproduction. The characteristic of minimal gene set can eliminate the possibility that other *Spiroplasma*-specific proteins are involved in the motility. Helicity switching was reconstituted by specific MreB pairs, MreB5−4, 5−1, 2−4, and 2−1 [14]. MreB is a bacterial actin homolog generally acting as a scaffold for cell wall synthesis and is not related to motility in most bacteria [15]. MreB5 labeling was succeeded by fusion with mCherry immediately after Y218 out of the total 350 residues, with retaining motility. The fluorescence was observed through the entire cell length [14].

Focusing on eukaryotic actins, they drive motility by two mechanisms [16]. One is as myosin rails, and the other is filament elongation and shortening coupled with regulatory proteins such as Arp2/3 complex. *Spiroplasma* motility is distinct from them because it is caused by only two MreB isoforms without actin regulatory proteins. This force-generating mechanism may be a hint to understand the divergency and evolution of molecular motors. Moreover, the mechanism may be applicable to actuators of “nanorobot” based on artificial liposomes.

In this study, we manipulated the expression level and fluorescence labeling, and then examined the functional differences between two MreBs. Moreover, we investigated MreB5 dynamics by using photobleaching and photoactivation analyses and a polymerization inhibitor.

## Materials and Methods

### Bacterial Strains and Culture Conditions

*S. eriocheiris* (TDA-040725-5T) and JCVI-syn3B (GenBank, CP069345.1) were cultured in SP4 medium [17] at 30℃ and 37℃, respectively, until they reached an optical density of 0.02−0.03 at 620 nm. For DNA manipulation, *Escherichia coli* strains DH5α and iVEC3 were used.

### Gene Manipulation of syn3B

DNA manipulation and JCVI-syn3B transformation were performed as previously described [18-20]. Plasmids and primers used in this study are listed in Figure S3 and Table S1, respectively. pSeN540 and pSeN540mC5 were prepared from pSeW545 and pSeW545-F5 (1), respectively by replacing the Ptuf promoter with the native promoter of MreB4. The tetracycline-inducible module was introduced from pSD079. A codon optimized PAmCherry gene was synthesized by the GenTitan Gene Fragment service (Genscript, NJ, US).For DNA assembly In-Fusion® HD Cloning Kit (Takara Bio Inc. Kusatsu, Japan), NEBuilder® HiFi DNA Assembly Master Mix (New England Biolabs, Inc., MA, USA) [20], and *Escherichia coli* (iVEC3) were used (Table. S1).

### Cell Treatments and Optical Microscopy

The following methods were used unless otherwise described. The cultured cells were centrifuged at 8,000×g for 5 min and suspended with supernatant at 5-fold cell density, if the cell density is not enough. The cell suspension was mixed in a ratio of 3 to 2 with a solution consisting of 20 mM HEPES (4-(2-hydroxyethyl)-1-piperazineethanesulfonic acid), 330 mM NaCl, 0.8% Methylcellulose, and flowed into the tunnel chamber [21] assembled with coverslips cleaned by KOH saturated ethanol. After 5−10 min, floating cells were removed by a flow of 30 μl solution consisting of 0.4% Methylcellulose, 20 mM HEPES, 330 mM NaCl. A22 (Cayman Chemical Company, Michigan, US) was dissolved in DMSO to be 10 mM. For A22 treatment, the liquid in a tunnel was replaced with a solution consisting of 250 μM A22, 20 mM HEPES, 330 mM NaCl, 0.4% Methylcellulose. Cells were observed by an inverted microscope IX71 (Evident, Tokyo, Japan) with a UPlanSApo 100×/1.4 NA Ph3 and CMOS camera (DMK33UX174, The Imaging Source, Bremen, Germany). Images were captured at 30 FPS (frame per second) and analyzed by ImageJ 1.54g.

For the identification of moving parts, the minimum intensity projection (Image > Stack > Z projection) was subtracted from each frame. For the comparison of promoter, the cultured cells were centrifuged at 9,000×g for 8 min and suspended with a phosphate-buffered saline (PBS) composed of 75mM sodium phosphate (pH7.3), 68mM NaCl, 20mM sucrose, at 20-fold cell density for observation.

### Protein Profile

For the expression induction, Tetracycline dissolved in ethanol at 4 mg/ml was added to the culture at a final concentration of 0.4 μg/ml. The culture was then incubated at 37℃ for various times, centrifuged at 13,200 ×g for 5 min, repeated twice and resuspended in 1×PBS at 50-fold cell density, lysed with 1/3 volume of SDS buffer (10% SDS, 20% Glycerol, 0.25M Tris-HCl pH6.8, 0.1% Bromophenol Blue, 20% β-mercaptoethanol) and ultrasonic treatment. The lysate was heated at 95°C for 3 min and subjected to SDS-10% or 12.5% PAGE. Loading volumes were normalized by culture ODs.

### Photobleaching and photoactivation

For FRAP (Fluorescence Recovery After Photobleaching), cells were observed by an inverted microscope IX83 with a UPlanSApo 100×/1.45 NA Ph3 (Evident) and an EMCCD camera (ixon897, Andor, Belfast, UK). Images were captured at 40 FPS. Fluorescence imaging was done with 10% of the maximum intensity of LED light source (U-LGPS, Evident) and manually switched to the maximum intensity with the minimum aperture during bleaching. Timelapse images were captured at 1 FPS with 1/30 s exposure. For chemical fixation, 4−5m strain was mixed in a ratio of 1 to 1 with an 8% paraformaldehyde solution, incubated at 22−27℃ for 10 min, rinsed with 1 M glycine. Fixed cells were centrifuged at 12,000×g for 10 min at 10°C and suspended with a solution consisting of 20 mM HEPES, 330 mM NaCl to be 2-fold cell density of the culture. Due to the small size of JCVI-syn3B cell, a circular bleached area was impossible unlike common way in FRAP. Additionally, tracking fluorescence intensity at a specific region was challenging due to cell movements. Therefore, edge parts of cells adhered linearly to the glass surface was bleached, and pixel values were profiled vertical to the cell axis, with width of 5 pixels. The fluorescence intensity of a frame was quantified by subtracting the minimum value from the maximum value in the selected regions.

For photoactivation, we irradiated 352-402 nm light, by a pattern illuminator (LEOPARD2-White, OPTO-LINE Saitama, Japan). In this setup, cells were observed by IX73 equipped with an sCMOS camera (Zyla4.2, Andor, UK). mCherry and PAmCherry signals were detected by using IX3-FMCHEXL (Ex.565-585 nm, Dichroic.595 nm, Em.600-690) filter (Evident).

## Results and Discussion

### New Constructs

To improve MreB expression levels, we replaced the Ptuf and Pspi promoters by the native promoter of MreB4 (PmreB4) from the *S. eriocheiris* (Fig. S4). In these strains, the expression of MreB4 and 5 was 2.6 ± 0.4- and 9.3 ± 1.1-fold higher than those in the previous strain, respectively (Fig. S1a). The ratio of moving cells increased from 17.5% to 50.4% (Movie. S1). Previous strains expressing only MreB4 or MreB5 under the Pspi promoter did not show helical shape or movements [14]. However, new strains under PmreB4 promoter resulted in non-moving but helical cells with some ratio (Fig. S1b). The protein amounts estimated from band intensity were 43 ± 7 and 14 ± 3-fold higher than the previous strains for only MreB4 or MreB5, respectively (Fig. S1a).

### Localization of Fluorescently Labeled MreBs

To examine the roles of MreB5 in elongation and motility of a cell, we observed the localization of mCherry-labeled MreB5 (MreB5m) in various forms of cells. A partial signal was found in almost all the cells, localizing at moving helical parts (Fig. S2, Fig. 1 left), indicating that MreB5 is involved in both helix formation and movements.

**Figure 1.**
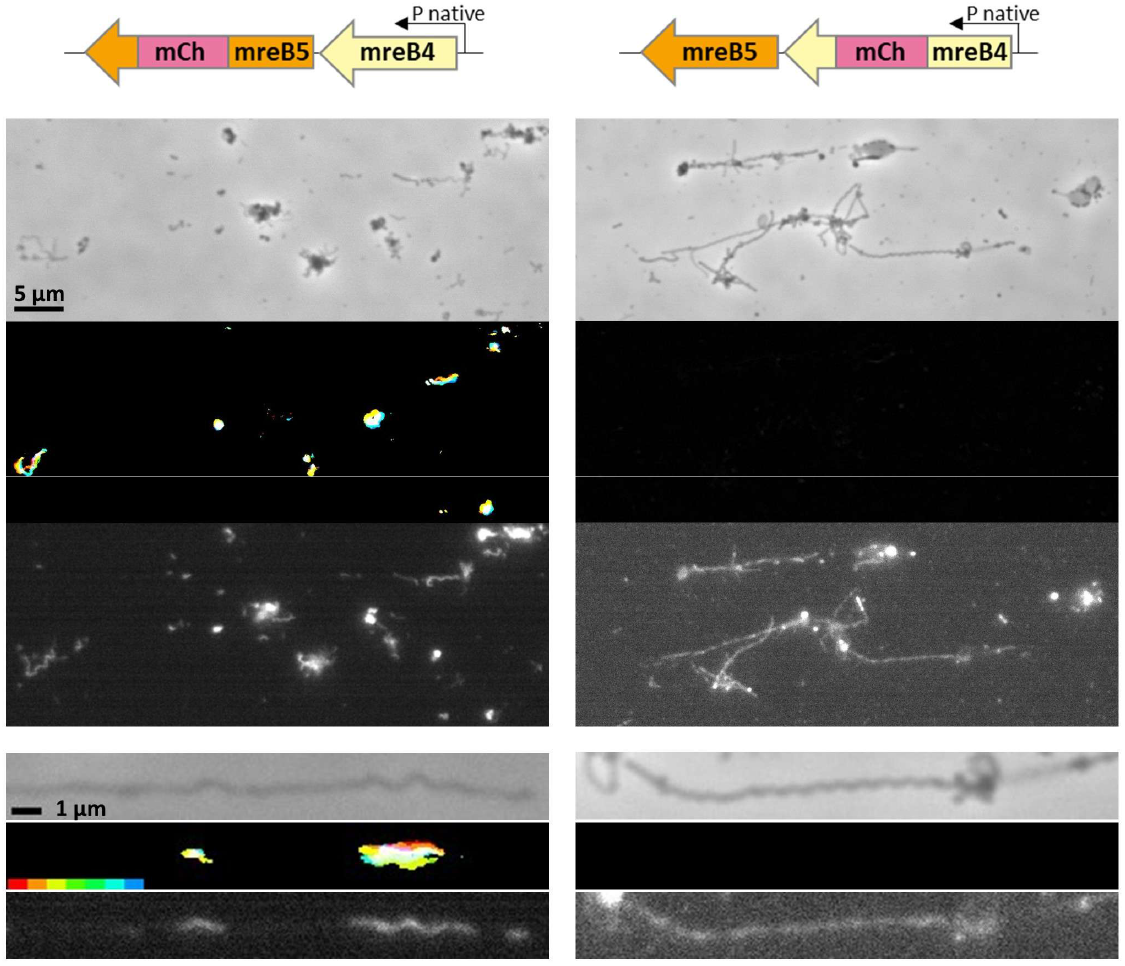
Cellular localization of each MreB fused with mCherry. Gene constructs are illustrated on the top. Field and cell images are shown for phase contrast, rainbow trace, fluorescence, from the top. In the rainbow trace, cell images with 1/30 s interval were colored from red to blue and overlaid. Excitation light was 10 times more intense for MreB4 than that for MreB5.

Next, to examine the subcellular localization of MreB4, we labeled it by inserting mCherry immediately after Y225, a position corresponding to the labeling site of MreB5. Signals were observed in almost all the cells, and some cells showed helical shape. However, no moving cells were observed (Fig. S3, Fig. 1 right). Then, we moved the insertion positions per one residue, however none of the 14 constructs showed cell movements (Fig. S3). We also tried to label MreB4 by small Tetracysteine (TC) motif composed of only 12 amino acid residues (FLNCCPGCCMEP). Biarsenical dye called FlAsH is expected to conjugate this motif and emit fluorescence [22,23]. We fused TC motif at N-terminus, C-terminus, and immediately after Y225 of MreB4. The construct inserted immediately after Y225 showed 5 cells moving out of approximately 500 (Movie. S2). However, the kink propagation was not observed, and the background fluorescence probably from unspecific binding of FlAsH was high. Then, we concluded that the TC labeling was not suitable for analysis of MreB cellular dynamics. The difficulty in fluorescence labeling of MreB4 may suggest that MreB4 interacts with other structures through larger numbers of positions than MreB5.

### Relation between Expression Level of each MreB and Motility

We examined the ratio of motile cells at various expression levels of each MreB. We constructed the strain with a constitutive expression of a MreB and another inducible one. In these strains, one MreB is under the control of the Pxyl/tetO2 promoter, where the expression is prevented by a tetracycline repressor (TetR) [24]. The MreB expression was induced by adding 9 µM tetracycline. The protein levels and ratio of motile cells were then analyzed (Fig. 2ab, Movie. S3). In the strain expressing both MreBs constitutively, the ratio of moving cell was 37.4±4.2%. In the strain expressing only MreB5m constitutively, motile cell ratio reached 37.6±3.4% when MreB4 was induced to 35.7±3.4% of the constitutive level. On the other hand, in the strain expressing only MreB4 constitutively, MreB5m induction to the 35.7±1.2% resulted in 8.6±2.9% motile cells. Higher requirement of MreB5 than MreB4 to move the cells may suggest that MreB5 serves as the backbone of the helix and that MreB4 causes MreB5 helix inversion. This may be related to the fact that MreB1, thought to function similarly as MreB4, has high ATPase activity [25]. However, it should be noted that the expression level shown here is for the whole culture and not for individual cells.

**Figure 2.**
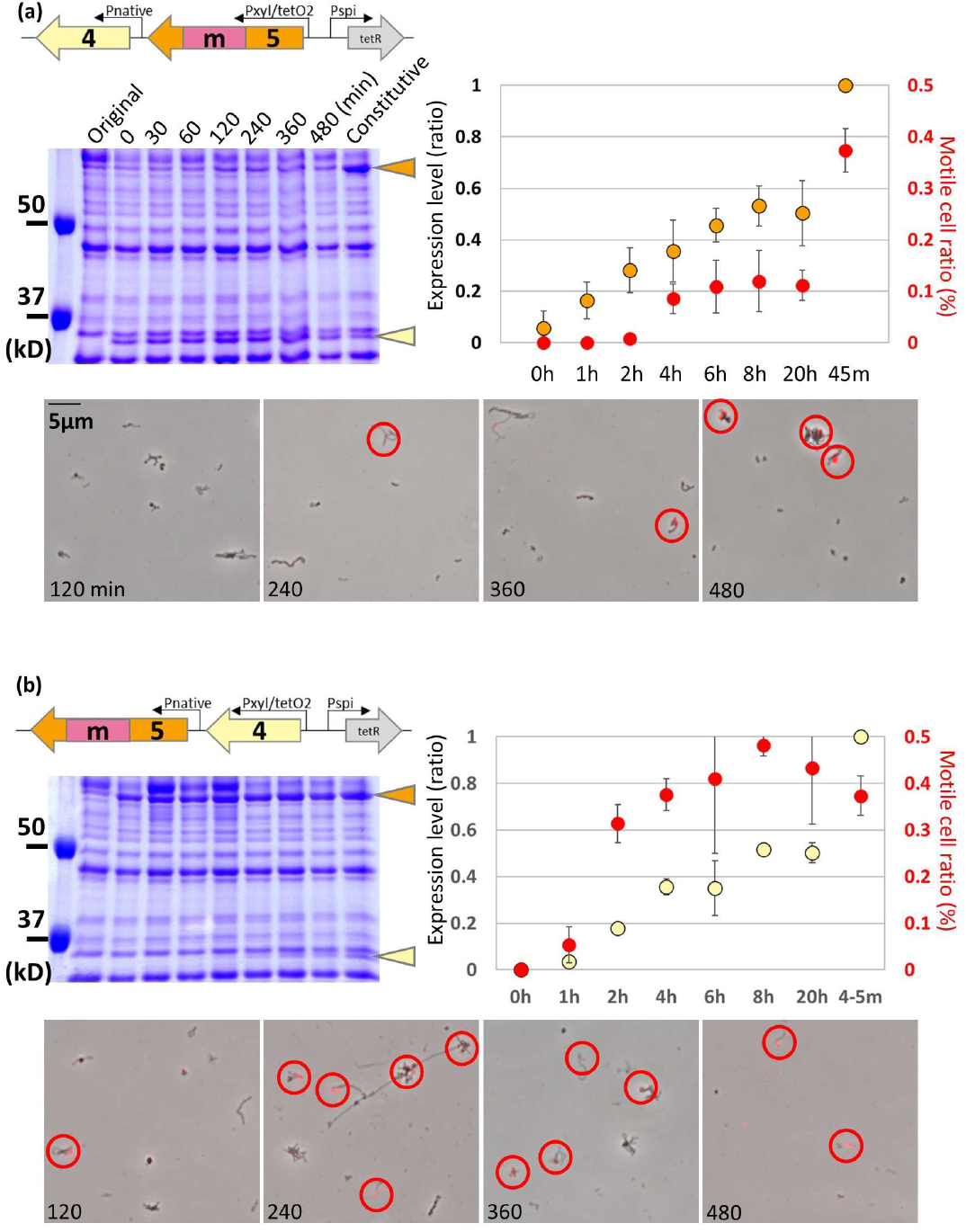
Motile cell ratio at various times after one MreB induction in a background of constitutive expression of another MreB. (a) MreB5 induction to constitutive MreB4 expression. (b) MreB4 induction to constitutive MreB5 expression. Gene constructs are illustrated in the upper left. (a, b) Protein profiles at various induction times were analyzed by SDS-10%-PAGE. The bands for MreB4 and MreB5m are shown by yellow and orange triangles, respectively. Cell images at various induction times are presented on the bottom. The moving parts are colored red. Motile cells are marked by a red circle. The integrated results (n=3) are shown in the upper right. Yellow and orange circles show the band intensity of MreB4 and MreB5m normalized by MreB5m of strain pSeN540mc5 expressing MreB5m constitutively. Red circles show the ratio of moving cells.

### MreB5 Replacement Traced by Photobleaching and Photoactivation

MreB generally has polymerization and ATPase activities like eukaryotic actin [16]. In *Spiroplasma*, MreB5 polymerizes into filaments in the presence of ATP [26]. To examine whether frequent polymerization-depolymerization is involved in the helicity switching, we focused on MreB5m behaviors in a moving cell. The FRAP is a technique to estimate the diffusion rate of target proteins from the recovery rate of the fluorescence intensity at the photobleached area [27]. If MreB5 subunit replacement is frequent, fluorescence should recover rapidly. Here, we focused on cells that were attached to glass only at poles and stretched by liquid flow applied before observation. These cells exhibited movements, although the kink propagation did not fully reach the pole due to the attachment. We bleached one end of such cells and traced the fluorescence intensities as shown by a kymograph (Fig. 3a). In the strain expressing MreB4 and MreB5m (Fig. 3a upper, Movie. S4), fluorescence intensity dropped to about 10% of initial value and returned to a plateau at about 30% within 5 s. In chemically fixed cells (Fig. 3a bottom), the recovery was not observed. In the strain expressing free mCherry additionally to MreB4 and MreB5 (Fig. 3a middle), the post-bleach intensity recovered to about 50% within 5 s. The latter two results proved that the fluorescence recovery in this experiment reflects the fluidity of the protein. The recovery to 30% in the strain expressing MreB4 and MreB5m indicates the presence of highly fluid MreB5m. However, the lack of subsequent recovery suggests that some MreB5m have low fluidity. To trace the behaviors of MreB forming filaments, we tried photoactivation using PAmCherry (Photoactivatable mCherry) [28]. This protein is initially non-fluorescent and activated by UV irradiation. The photo switching is caused by only 10 amino acid modification from mCherry. The MreB5 fused with PAmCherry retained functions for cell elongation and movements, as observed for mCherry fusion. Then, we irradiated the cells with UV light, and traced the subsequent fluorescence intensity (Fig. 3b, Movie. S5). After UV irradiation, only the irradiated area emitted fluorescence. The fluorescence intensity decreased gradually due to the photobleaching, however there was no obvious inflow or outflow of MreB5m between bright and dark regions. These results indicate that MreB5 remains in the same cellular position during movements, without obvious replacements. This seems to be distinct from other bacterial actins such as ParM and MamK that exhibit dynamic polymerization [29,30].

**Figure 3.**
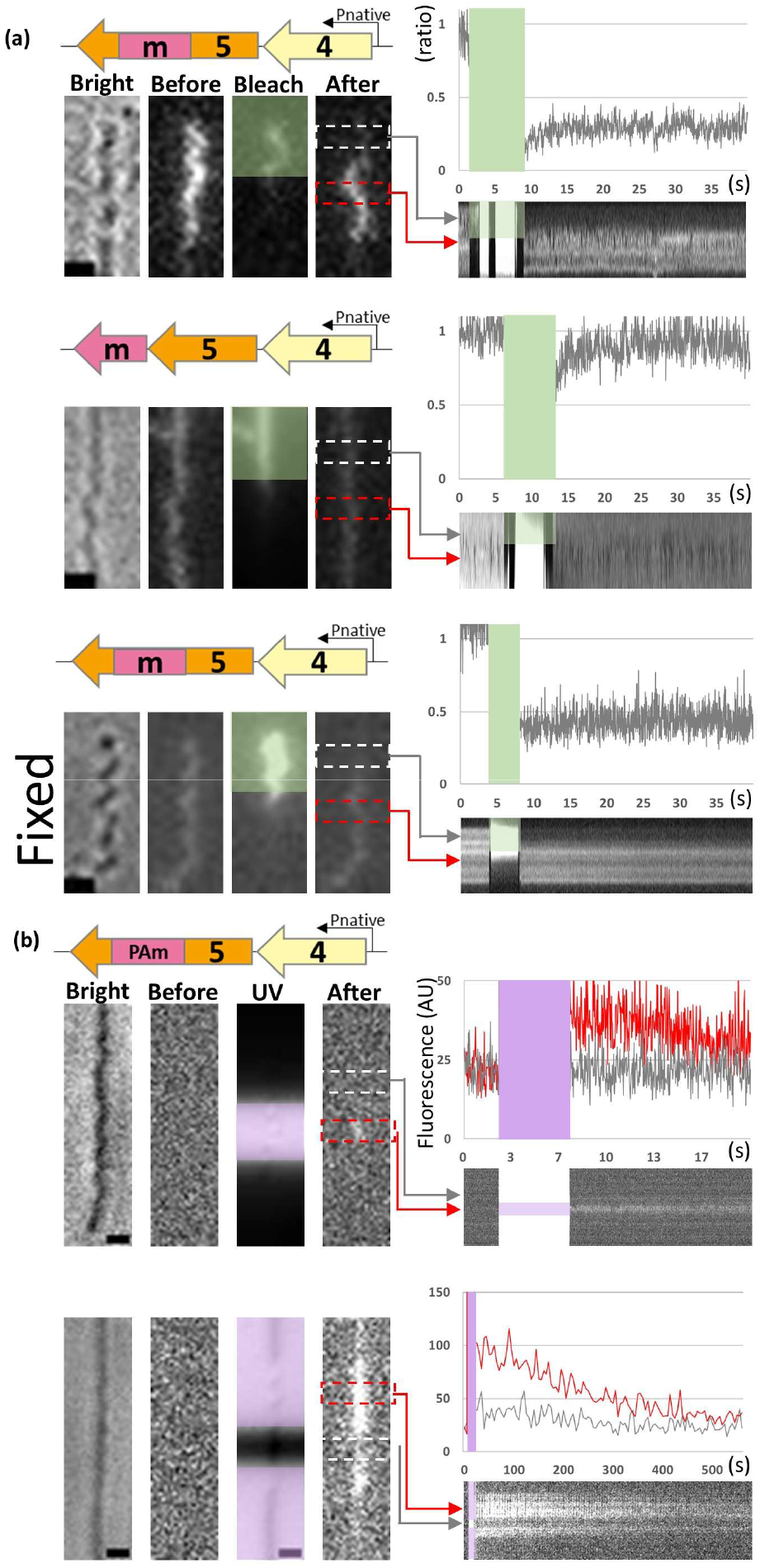
MreB5 dynamics examined by photobleaching and photoactivation. In each block, gene constructs are illustrated in the upper left. Cell images are shown in the left. Scale bar, 1μm. Fluorescence traces are shown in the right upper. Kymographs are shown in the right lower, (a) Photobleaching. Cell images in brightfield, prebleach, bleach, postbleach are shown. Bleached area is shown by a green translucent rectangle. Recovery of bleached fluorescence was traced for 40 s by measuring the fluorescence intensity ratio of white to red boxed areas. Bleaching time is shown by a green rectangle. (b) Photoactivation. Cell images in brightfield, preactivation, UV irradiation, postactivation are shown. UV irradiated area is shown by a purple translucent rectangle. Fluorescence intensity for red and white boxed areas are traced. UV irradiation time is shown by a purple rectangle.

### Effect of MreB Polymerization Inhibitor A22 on Cell Movements and MreB5 Localization

To know the role of the polymerization ability of MreB, we examine the effects of an MreB polymerization inhibitor A22. A22 is a compound with a molecular weight of 271.6 and has been shown to bind the nucleotide binding site of *Caulobacter crescentus* MreB [31,32]. Addition of 250 µM of A22 to the syn3B cells expressing MreB4 and MreB5m linearized the helix, and stalled movements in 150 s (Fig. 4a, Movie. S6). To confirm if the intracellular MreB5 become diffusive, we added A22 after the photobleaching in FRAP observation. Fluorescence that once reached a plateau recovered by 20% after A22 addition (n=2)(Fig. 4b). For the same purpose, we added A22 to the cell expressing MreB4 and MreB5-PAmCherry after the UV irradiation. Fluorescence of the specific photoactivated region was observed to be spread in 100 s after A22 addition (Fig. 4c). These results indicate that A22 inhibited some interaction responsible for MreB positioning in a cell. Next, we initially linearized the cell by A22, then performed FRAP observation. If the MreB5 is released from all interactions, fluorescence trace is expected to be like free mCherry (Fig. 3a middle). However, no fluorescence recovery was observed (Fig. 4d). This result may suggest that some MreB5m molecules stay at their positions even after the cell linearization. This may be consistent with the uneven PAmCherry fluorescence distribution after A22 addition (Fig. 4c).

**Figure 4.**
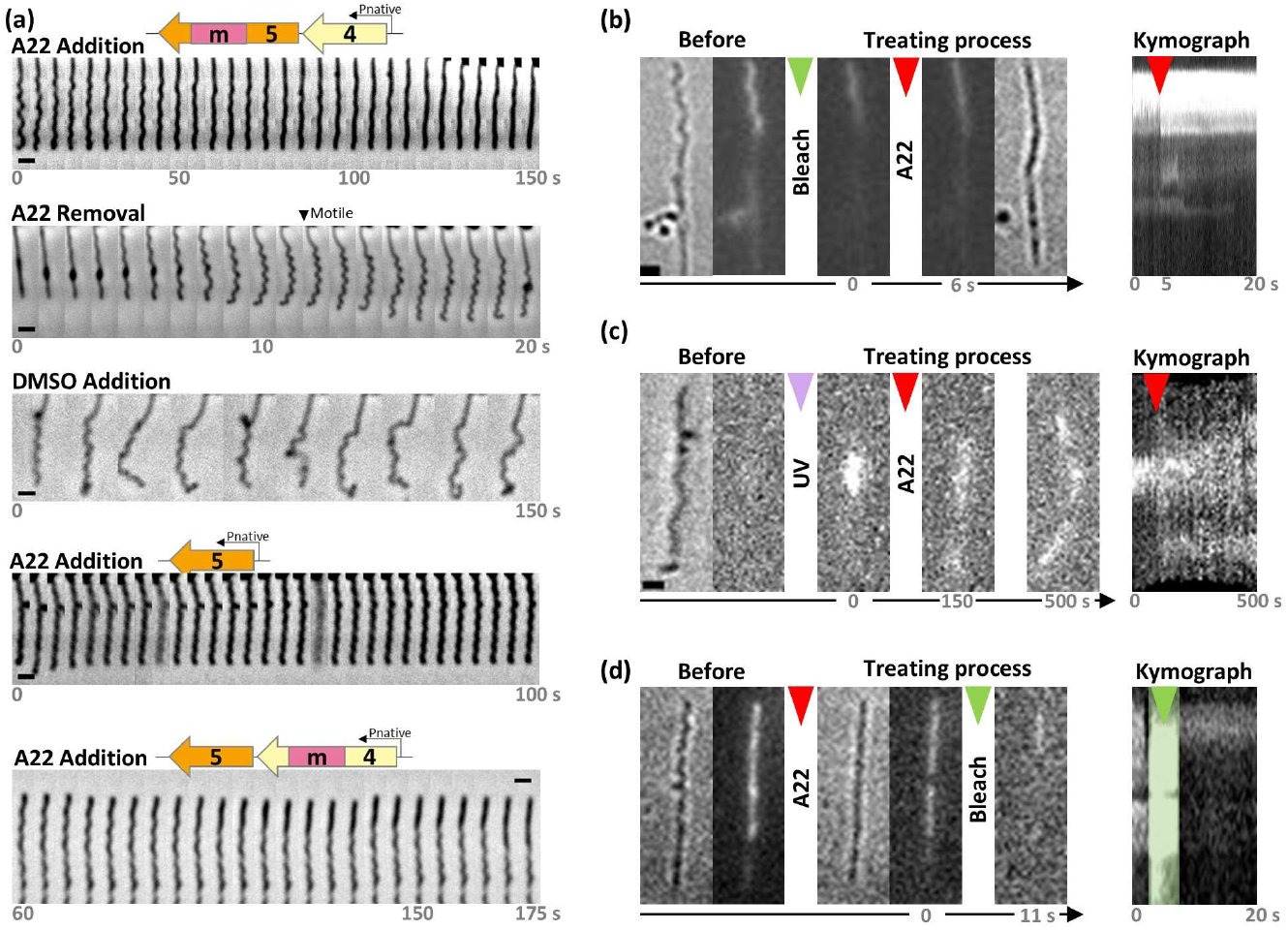
Effects of A22 on cell movements, shape, and MreB5m behaviors. **(a)** Change in cell shape after addition and removal of A22. DMSO was used for control. Gene constructs are illustrated on the top. **(b)-(d)** Effects of A22 addition on MreB5 behaviors. Cell images in the process are shown in order. Kymograph of fluorescence distribution along the cell axis is shown in the right. **(b)** Cells were first photobleached (green triangle) and treated with A22 (red triangle). The kymograph starts after the photobleaching. **(c)** Cells were first photoactivated (purple triangle) and treated with A22 (red triangle). The kymograph starts after the photoactivation. **(d)** Cells were first treated with A22 (red triangle) and photobleached (green triangle). The kymograph starts after the A22 addition.

Besides the above experiments, we observed the cell changes after the A22 removal to know if this motility inhibition is reversible or not. A22 removal caused helix reformation immediately and the motility recovered within 10 s (Fig. 4a, Movie. S7). Since both helix linearization and reformation occurred along the entire cell axis, MreB5 unlikely were released only at cell poles.

We added A22 to the immotile cells. Cells expressing intact MreB5 without MreB4 were hardly linearized by A22 (Fig. 4a, Movie. S8). We also added A22 to a cell with intact MreB5 and labeled MreB4. As a result, only 40% was linearized after 110−175 s from A22 addition (Fig. 4a). This observation can be explained by two possibilities. One is that the A22 binding site of MreB5 is not accessible in the absence of movements. Another possibility is that A22 binds to MreB protein, but the conformational change does not occur.

## Conclusion

This study suggests (i) Two MreBs have distinct functions, (ii) MreB5 does not have a dynamic replacement as observed in treadmilling of eukaryotic actin, (iii) Helicity switching is linked to conformational change in MreB5 (Fig. 5).

**Figure 5.**
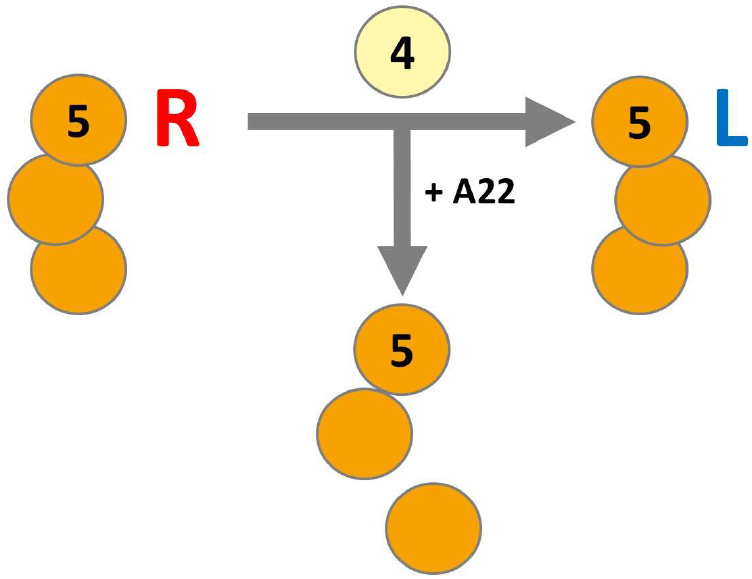
Summary of this study. Right-handed structure of MreB5 turns to Left-handed without obvious subunit replacements when MreB4 exists. A22 addition to moving cell leads to the MreB5 spreading.

## Author Contributions

YT performed most of the experiments and wrote the original draft. HK performed the experiment described in the section “New constructs”, and provided the strategy of inducible MreB constructs. YVM provided the photoactivation experiment with equipments. YT and MM edited the manuscript. All authors contributed to the project and manuscript.

## Data Availability

The evidence data generated/analyzed in this study are included in this article.

A preliminary version of this work, DOI: https://doi.org/10.1101/2025.05.23.655722, was deposited in the bioRxiv on May 25, 2025.

## Acknowledgements

This study was supported by JST CREST grant (grant number JPMJCR19S5 to MM), and Grant-in-Aid for JSPS Fellows (grant number JP24KJ0189 to HK).

---

**Supplementary Figure S1.**
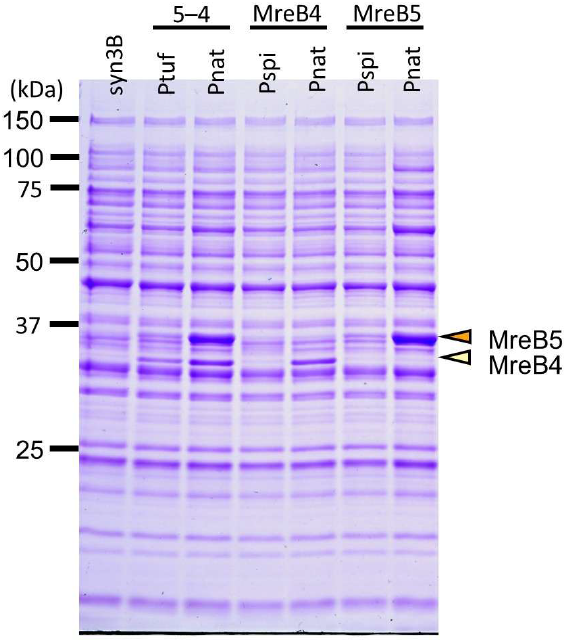
Comparison of expression level of MreB4 and MreB5 between previous and improved promoters, by SDS-12.5% PAGE. The bands for MreB4 and MreB5 are marked by yellow and orange triangles, respectively.

**Supplementary Figure S2.**
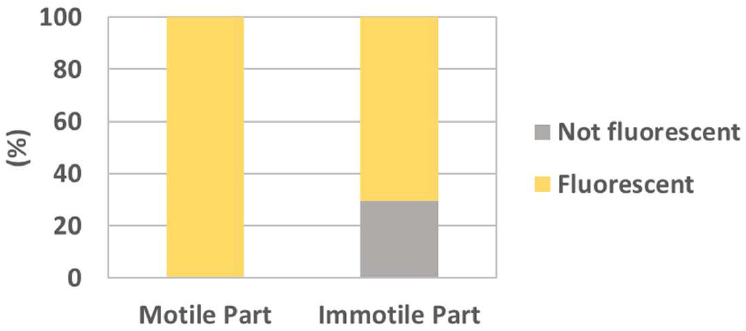
The ratio of fluorescent parts at moving and non-moving parts of cells harboring pSeN540mc5.

**Supplementary Figure S3.**
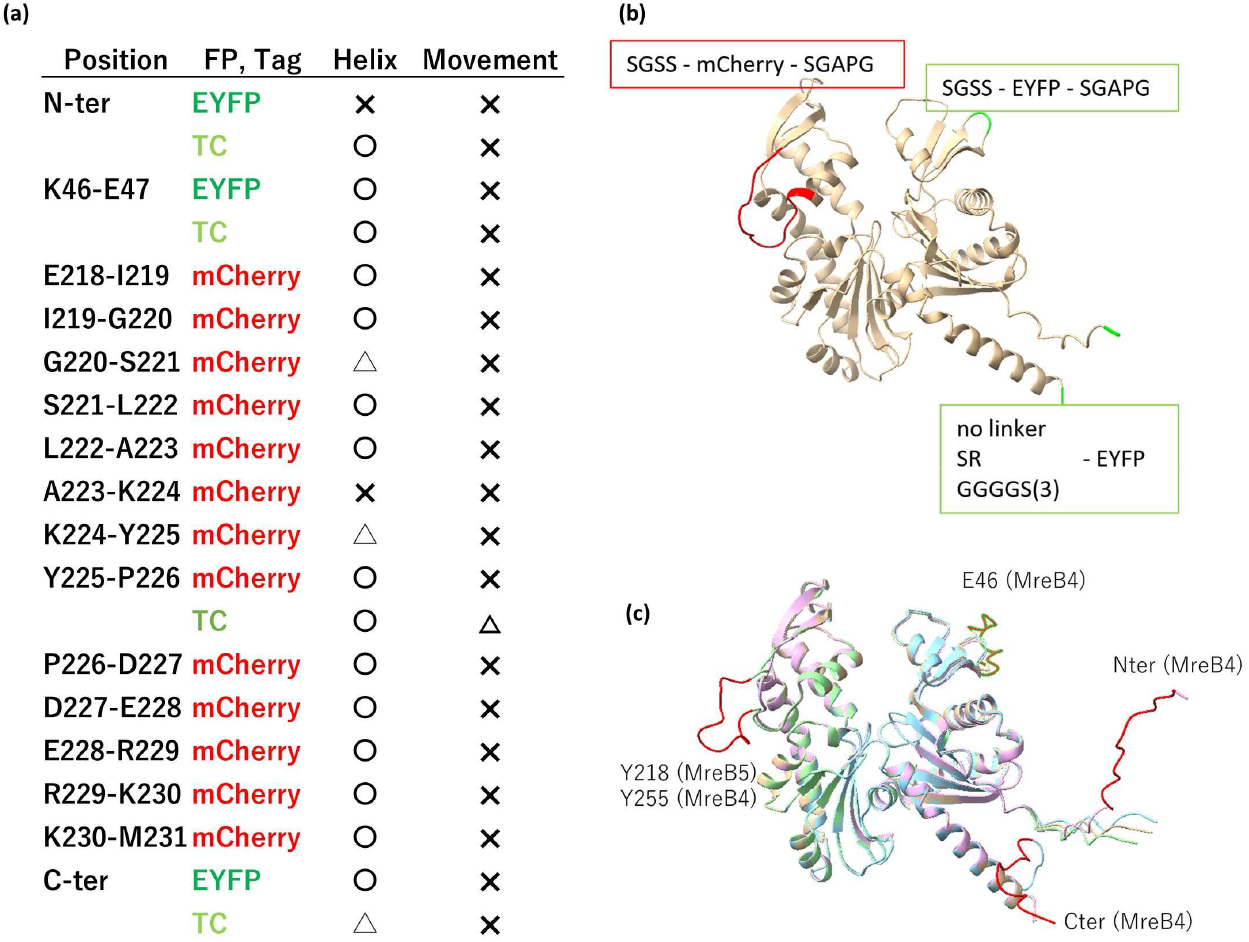
Characterizing transformants of MreB4 fused with mCherry, EYFP, and TC-tag. (a) Morphology and movements. (b) Labeled positions indicated in MreB4 structure predicted by AlphaFold2. (c) Predicted structure of MreB4 labeled with TC-Tag.

**Supplementary Figure S4.**
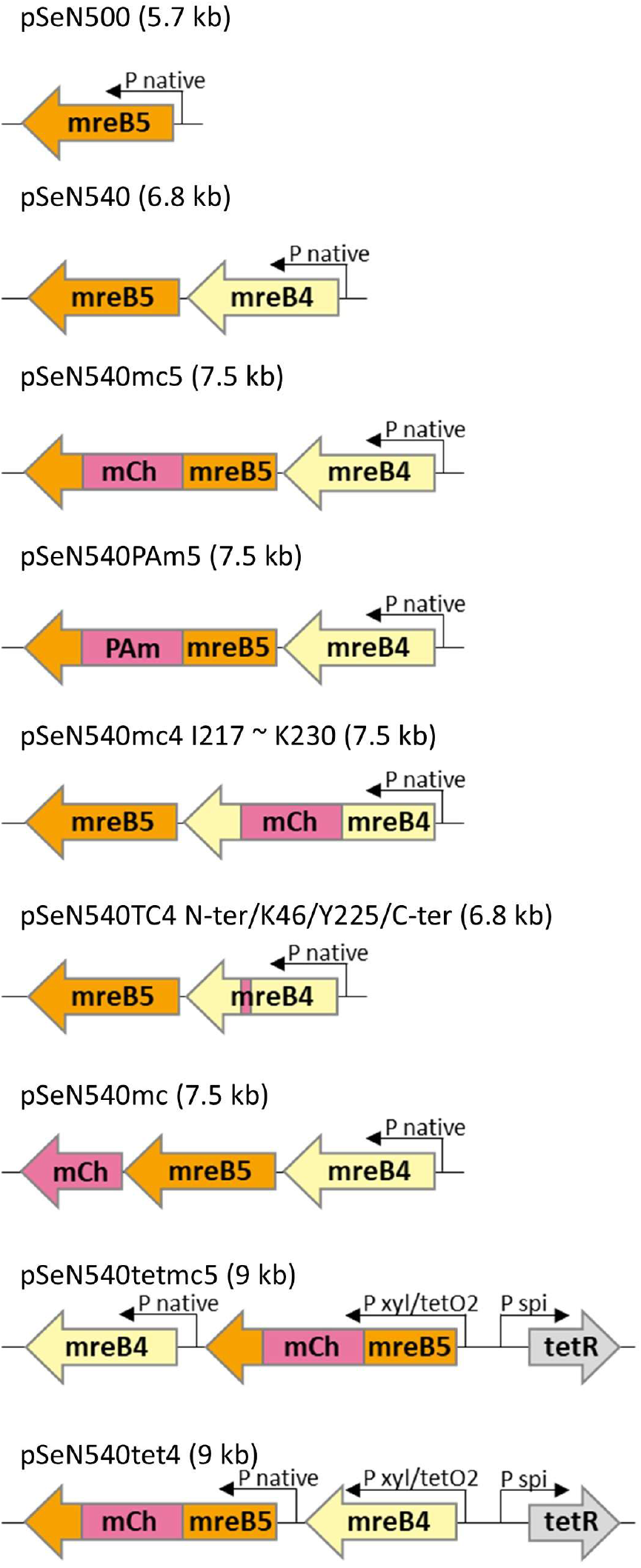
Schematic presentation for DNA constructs used in this study.

**Supplementary Table S1.**
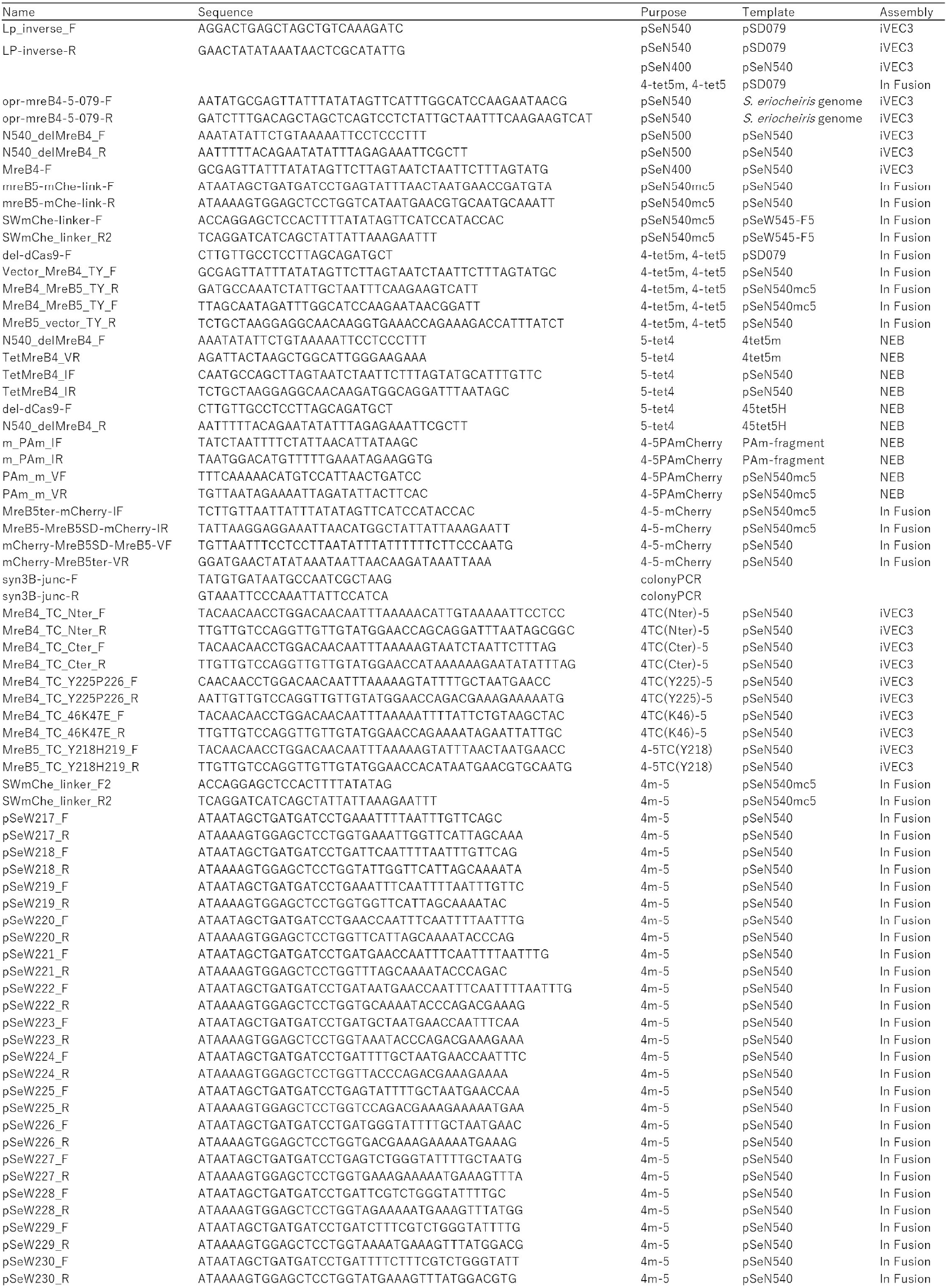
DNA primers used in this study.

## References

[1] Harne, S., Gayathri, P., Beven, L. Exploring Spiroplasma biology: Opportunities and challenges. Front. Microbiol. 10, 589279 (2020). 10.3389/fmicb.2020.589279

[2] Kakizawa, S., Mizutani, M. Overview of the new genus names for Mycoplasmas. Japanese Society of Microbial Ecology 40, 29–37 (2025). 10.20709/jsmeja.40.1_29

[3] Wang, Y., Huang, J.M., Zhou, Y.L., Almeida, A., Finn, R.D., Danchin, A., et al. Phylogenomics of expanding uncultured environmental Tenericutes provides insights into their pathogenicity and evolutionary relationship with Bacilli. BMC Genomics 21, 408 (2020). 10.1186/s12864-020-06807-4

[4] Shaevitz, J.W., Lee, J.Y., Fletcher, D.A. Spiroplasma swim by a processive change in body helicity. Cell 122, 941–945 (2005). 10.1016/j.cell.2005.07.004

[5] Kurner, J., Frangakis, A.S., Baumeister, W. Cryo-electron tomography reveals the cytoskeletal structure of Spiroplasma melliferum. Science 307, 436–438 (2005). 10.1126/science.1104031

[6] Nakane, D., Ito, T., Nishizaka, T. Coexistence of two chiral helices produces Kink translation in Spiroplasma swimming. J. Bacteriol. 202 (2020). 10.1128/JB.00735-19

[7] Liu, P., Zheng, H., Meng, Q., Terahara, N., Gu, W., Wang, S., et al. Chemotaxis without conventional two-component system, based on cell polarity and aerobic conditions in helicity-switching swimming of Spiroplasma eriocheiris. Front. Microbiol. 8, 58 (2017). 10.3389/fmicb.2017.00058

[8] Sasajima, Y., Kato, T., Miyata, T., Kawamoto, A., Namba, K., Miyata, M. Isolation and structure of the fibril protein, a major component of the internal ribbon for Spiroplasma swimming. Front. Microbiol. 13, 1004601 (2022). 10.3389/fmicb.2022.1004601

[9] Trachtenberg, S., Dorward, L.M., Speransky, V.V., Jaffe, H., Andrews, S.B., Leapman, R.D. Structure of the cytoskeleton of Spiroplasma melliferum BC3 and its interactions with the cell membrane. J. Mol. Biol. 378, 778–789 (2008). 10.1016/j.jmb.2008.02.020

[10] Williamson, D.L., Renaudin, J., Bove, J.M. Nucleotide sequence of the Spiroplasma citri fibril protein gene. J. Bacteriol. 173, 4353–4362 (1991). 10.1128/jb.173.14.4353-4362.1991

[11] Ku, C., Lo, W.S., Kuo, C.H. Molecular evolution of the actin-like MreB protein gene family in wall-less bacteria. Biochem. Biophys. Res. Commun. 446, 927–932 (2014). 10.1016/j.bbrc.2014.03.039

[12] Hutchison, C.A., 3rd, Chuang, R.Y., Noskov, V.N., Assad-Garcia, N., Deerinck, T.J., Ellisman, M.H., et al. Design and synthesis of a minimal bacterial genome. Science 351, aad6253 (2016). 10.1126/science.aad6253

[13] Pelletier, J.F., Sun, L., Wise, K.S., Assad-Garcia, N., Karas, B.J., Deerinck, T.J., et al. Genetic requirements for cell division in a genomically minimal cell. Cell 184, 2430–2440 e2416 (2021). 10.1016/j.cell.2021.03.008

[14] Kiyama, H., Kakizawa, S., Sasajima, Y., Tahara, Y.O., Miyata, M. Reconstitution of a minimal motility system based on Spiroplasma swimming by two bacterial actins in a synthetic minimal bacterium. Sci. Adv. 8, eabo7490 (2022). 10.1126/sciadv.abo7490

[15] Shi, H., Bratton, B.P., Gitai, Z., Huang, K.C. How to build a bacterial cell: MreB as the foreman of E. coli construction. Cell 172, 1294–1305 (2018). 10.1016/j.cell.2018.02.050

[16] van den Ent, F., Amos, L.A., Lowe, J. Prokaryotic origin of the actin cytoskeleton. Nature 413, 39–44 (2001). 10.1038/35092500

[17] Tully, J.G., Rose, D.L., Whitcomb, R.F., Wenzel, R.P. Enhanced isolation of Mycoplasma pneumoniae from throat washings with a newly-modified culture medium. J. Infect. Dis. 139, 478–482 (1979). 10.1093/infdis/139.4.478

[18] Karas, B.J., Wise, K.S., Sun, L., Venter, J.C., Glass, J.I., Hutchison, C.A., 3rd, et al. Rescue of mutant fitness defects using in vitro reconstituted designer transposons in Mycoplasma mycoides. Front. Microbiol. 5, 369 (2014). 10.3389/fmicb.2014.00369

[19] Mizutani, M., Glass, J.I., Fukatsu, T., Suzuki, Y., Kakizawa, S. Robust and highly efficient transformation method for a minimal mycoplasma cell. J. Bacteriol. 207, e0041524 (2025). 10.1128/jb.00415-24

[20] Uenoyama, A., Kiyama, H., Mimura, M., Miyata, M. Rapid in vitro method to assemble and transfer DNA fragments into the JCVI-syn3B minimal synthetic bacterial genome through Cre/loxP system. Biophys Physicobiol 21, e210024 (2024). 10.2142/biophysico.bppb-v21.0024

[21] Kasai, T., Hamaguchi, T., Miyata, M. Gliding motility of Mycoplasma mobile on uniform oligosaccharides. J. Bacteriol. 197, 2952–2957 (2015). 10.1128/JB.00335-15

[22] Griffin, B.A., Adams, S.R., Tsien, R.Y. Specific covalent labeling of recombinant protein molecules inside live cells. Science 281, 269–272 (1998). 10.1126/science.281.5374.269

[23] Martin, B.R., Giepmans, B.N., Adams, S.R., Tsien, R.Y. Mammalian cell-based optimization of the biarsenical-binding tetracysteine motif for improved fluorescence and affinity. Nat. Biotechnol. 23, 1308–1314 (2005). 10.1038/nbt1136

[24] Mariscal, A.M., Kakizawa, S., Hsu, J.Y., Tanaka, K., Gonzalez-Gonzalez, L., Broto, A., et al. Tuning gene activity by inducible and targeted regulation of gene expression in Minimal bacterial cells. ACS Synth. Biol. 7, 1538–1552 (2018). 10.1021/acssynbio.8b00028

[25] Takahashi, D., Kiyama, H., Matsubayashi, H.T., Fujiwara, I., Miyata, M. A bacterial actin with high ATPase activity regulates the polymerization of a partner MreB isoform essential for Spiroplasma swimming motility. J. Biol. Chem. 301, 110462 (2025). 10.1016/j.jbc.2025.110462

[26] Takahashi, D., Fujiwara, I., Miyata, M. Purification and ATPase activity measurement of Spiroplasma MreB. Methods Mol. Biol. 2646, 359–371 (2023). 10.1007/978-1-0716-3060-0_30

[27] Esue, O., Rupprecht, L., Sun, S.X., Wirtz, D. Dynamics of the bacterial intermediate filament crescentin in vitro and in vivo. PLoS One 5, e8855 (2010). 10.1371/journal.pone.0008855

[28] Subach, F.V., Patterson, G.H., Manley, S., Gillette, J.M., Lippincott-Schwartz, J., Verkhusha, V.V. Photoactivatable mCherry for high-resolution two-color fluorescence microscopy. Nat. Methods 6, 153–159 (2009). 10.1038/nmeth.1298

[29] Popp, D., Robinson, R.C. Many ways to build an actin filament. Mol. Microbiol. 80, 300–308 (2011). 10.1111/j.1365-2958.2011.07599.x

[30] Charles-Orszag, A., Petek-Seoane, N.A., Mullins, R.D. Archaeal actins and the origin of a multi-functional cytoskeleton. J. Bacteriol. 206, e0034823 (2024). 10.1128/jb.00348-23

[31] Iwai, N., Nagai, K., Wachi, M. Novel S-benzylisothiourea compound that induces spherical cells in Escherichia coli probably by acting on a rod-shape-determining protein(s) other than penicillin-binding protein 2. Biosci. Biotechnol. Biochem. 66, 2658–2662 (2002). 10.1271/bbb.66.2658

[32] van den Ent, F., Izore, T., Bharat, T.A., Johnson, C.M., Lowe, J. Bacterial actin MreB forms antiparallel double filaments. Elife 3, e02634 (2014). 10.7554/eLife.02634

